# Septins and cytokinesis in the polymorphic fungus *Aureobasidium pullulans*

**DOI:** 10.64898/2026.01.20.700616

**Authors:** Analeigha V. Colarusso, Alison C.E. Wirshing, Daniel J. Lew

## Abstract

During cytokinesis of animals and fungi, a contractile actomyosin ring (CAR) assembles at target locations and constricts to drive cell separation. In animal cells, the position of the CAR is determined by the mitotic spindle, so that the cleavage plane is perpendicular to the mitotic axis. However, in budding yeasts, the location of CAR assembly is specified by a cortical septin cytoskeleton that recruits CAR components to the neck. In the polymorphic fungus *Aureobasidium pullulans*, we show that septins assemble at mother-bud necks and predict the site of CAR assembly. Cells lacking septins stochastically failed to assemble CARs at a subset of bud necks. However, even cells lacking all four core septins were able to assemble CARs at 75% of bud necks. Our findings suggest the existence of a novel CAR positioning strategy that requires neither septin scaffolds nor nuclear/spindle cues to enable CAR assembly and constriction at bud necks.

**eTOC SUMMARY:** Budding yeasts are thought to use septins to mark mother-bud necks as sites for cytokinesis. Here, we find that the multibudding yeast *Aureobasidium pullulans* can position cytokinetic machinery at most bud necks even in the absence of septins, revealing a novel pathway to mark cytokinesis sites.

## INTRODUCTION

Animal and fungal cells use a contractile actomyosin ring (CAR) to constrict the cleavage furrow during cytokinesis (Glotzer, 2017). Orthogonal placement of the CAR relative to the mitotic spindle ensures that cell division endows each daughter with at least one copy of the full genome, but the detailed mechanisms used to position the CAR are quite diverse (Oliferenko et al., 2009). In most animal cells, cues from the mitotic spindle instruct the position of CAR assembly (Fig. 1A). However, there are exceptions. In the early embryo of the fruit fly *Drosophila melanogaster*, formation of primordial germ cells proceeds via membrane budding (Chen et al., 2025), and placement and constriction of the CAR at the “neck” of the bud occurs independent of the mitotic spindle (Cinalli and Lehmann, 2013). The CAR positioning mechanism in this case is unknown.

**Figure 1:**
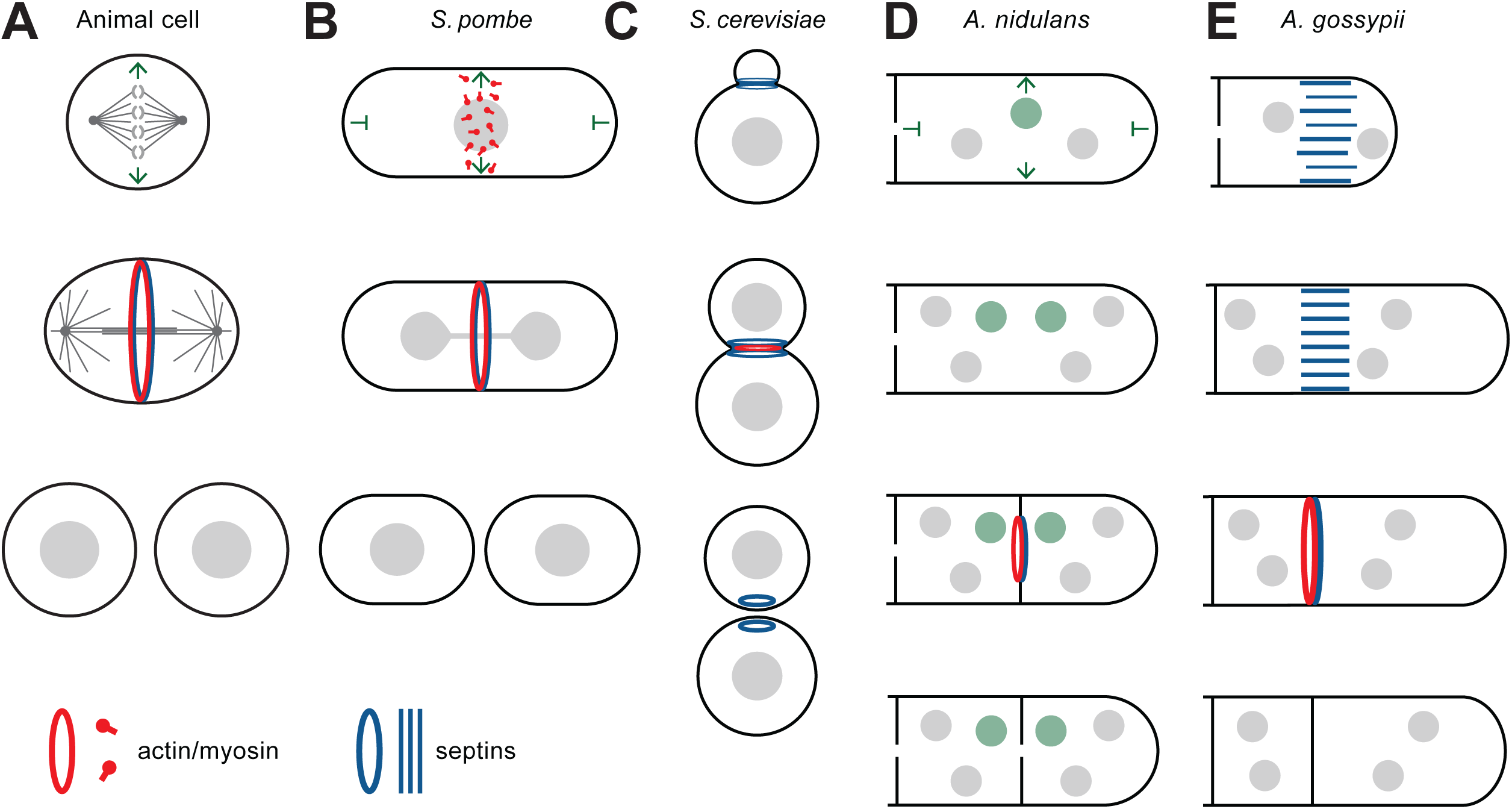
Defining the cytokinesis site. Proposed mechanisms to determine the location of contractile actomyosin rings (CARs) in different systems. (A) Animal cells: instructive cues (green arrows) from the mitotic spindle determine CAR (red) placement. (B) *S. pombe*: an instructive cue from the mitotic nucleus and inhibitory cues from cell poles determine CAR placement. In both animals and fission yeast, septins (blue) accumulate at the cleavage site but are not required for CAR placement. (C) *S. cerevisiae*: polarity cues place a septin scaffold (blue) that marks the bud neck and dictates CAR positioning. (D) *A. nidulans*: positive cues from a subset of mitotic nuclei (green) and inhibitory cues from hyphal tips/septa are thought to influence CAR placement. (E) *A. gossypii*: polarity cues position septin scaffolds that mark potential future CAR sites.

In fungi, the mechanisms of CAR positioning and assembly are best understood in the model yeasts *Schizosaccharomyces pombe* (fission yeast) and *Saccharomyces cerevisiae* (budding yeast), which use strikingly different strategies. In *S. pombe*, the CAR assembles from a set of precursor “nodes” (large protein complexes) at the cortex in the cell midzone. Positioning of the nodes occurs via somewhat redundant pathways that include node-inhibitory factors acting at cell tips (Celton-Morizur et al., 2006; Padte et al., 2006; Huang et al., 2007), and the positively acting factor Mid1, related to animal anillin (Sohrmann et al., 1996; Paoletti and Chang, 2000; Motegi et al., 2004) that is released from the centrally positioned nucleus in mitosis (Bähler et al., 1998). Interestingly, Mid1 does not concentrate in the nuclei of the related fission yeast *Schizosaccharomyces japonicus*, where another as-yet uncharacterized factor links nuclear position to actin assembly for CAR formation (Gu and Oliferenko, 2015). Septins are recruited to the cytokinesis site shortly before CAR constriction in fission yeast, but they are not needed for CAR placement or constriction (Wu et al., 2003, 2010). As with the cues from the mitotic spindle in animal cells, positioning cues from the mitotic nucleus provide a direct mechanism to coordinate chromosome segregation with the placement of the cleavage plane (Fig. 1B).

A fundamentally different strategy is employed by budding yeasts like *S. cerevisiae* (Fig. 1C). Building a bud automatically defines a mother-bud neck, where cytokinesis will occur once bud formation is complete. Given that pre-defined cleavage plane, the mitotic spindle must be oriented so as to segregate the daughter chromosomes across the neck (Segal and Bloom, 2001). After mitosis, the CAR assembles at the neck, in a manner that does not depend on spatial information from the mitotic spindle (Howell and Lew, 2012; Bhavsar-Jog and Bi, 2017). To mark the neck as the site of future CAR assembly, *S. cerevisiae* polarity factors recruit and organize a collar of cortical septins at the emerging bud neck. The core septins Cdc3, Cdc10, Cdc11, and Cdc12 form octamers that polymerize into a meshwork of filaments, creating scaffold that remains stably localized at the neck and progressively recruits CAR proteins through the cell cycle, with actin arriving only after mitosis to assemble the full CAR (Fig. 1C) (Gladfelter et al., 2001; Ong et al., 2014; Bhavsar-Jog and Bi, 2017; Okada et al., 2021). In all budding yeasts where they have been examined, septins assemble a scaffold at the bud neck that is presumed to position the CAR (Böhmer et al., 2009; Sudbery, 2001; Warenda and Konopka, 2002; Kozubowski and Heitman, 2010; Rippert and Heinisch, 2016; Petrucco et al., 2024). However, it is unclear whether septins are universally required for neck-located cytokinesis, particularly in the distantly related basidiomycete budding yeasts (Kozubowski and Heitman, 2010; Altamirano et al., 2017).

Mechanisms of CAR positioning are less clear in other fungi. There are indications that pathways related to those in the model yeasts may apply quite broadly. Like *S. pombe* cells, hyphae are tubular in shape, and CARs construct septa perpendicular to the tube axis (Fig. 1D). In *Aspergillus nidulans*, nuclear positioning and microtubule spindle-derived cues appear to influence CAR positioning (Wolkow et al., 1996; Momany and Hamer, 1997). Unlike most yeasts, hyphae are multinucleate, and CARs only form near a subset of nuclei, usually those that are distant from both the hyphal tip and other septa. Thus, it may be that as in *S. pombe*, a combination of positive cues from nuclei/spindles and inhibitory cues from tips/septa act to position the CARs in *A. nidulans* (Seiler and Justa-Schuch, 2010) (Fig. 1D). In *Ashbya gossypii*, another hyphal species that is more closely related to *S. cerevisiae*, it appears that septins may be used to demarcate future septation sites (Seiler and Justa-Schuch, 2010) (Fig. 1E). In this system, septin structures are deposited at intervals by the tips of the growing hyphae and mark future septation sites (Helfer and Gladfelter, 2006; DeMay et al., 2009; Kaufmann and Philippsen, 2009). In contrast, septins are only recruited to sites of septation after mitosis and are not required to position septa in *A. nidulans* (Hernández-Rodríguez and Momany, 2012; Westfall and Momany, 2002).

Broadly speaking, the studies in fungi suggest that there are at least two fundamentally distinct ways to spatially coordinate spindle position and cytokinesis. In one strategy, spatial cues emerge from nuclei or mitotic spindles to position the CAR, assisted by inhibitory cues emanating from cell ends (Fig. 1B,D). In the other strategy, cortical sites are marked by stable septin scaffolds during morphogenesis to delineate the future CAR location, and nuclei or mitotic spindles then segregate the chromosomes across these predetermined cleavage sites (Fig. 1C,E).

Here, we analyzed septin behavior and function in the multibudding black yeast *Aureobasidium pullulans*. *A. pullulans* can grow as a budding yeast, and localizes septins to the bud neck (Petrucco et al., 2024) suggesting a septin-mediated CAR placement strategy. However, unlike most budding yeasts, *A. pullulans* mother cells are often multinucleate and able to generate multiple buds in the same cell cycle (Mitchison-Field et al., 2019; Petrucco et al., 2024; Wirshing et al., 2024). *A. pullulans* can also grow in a filamentous manner, generating branched hyphal networks, and make large multinucleate cells that divide medially (Cooke, 1959; Kocková-Kratochvílová et al., 1980; Ramos and García Acha, 1975; Seviour et al., 1984). This morphological plasticity provides an opportunity to investigate how mechanisms of CAR placement, assembly, and constriction might be adapted for cells that differ in size and shape, but share a common genome. We report that while septins improve the efficiency of CAR assembly, they are surprisingly not needed to specify the bud neck as a site for CAR positioning.

## RESULTS

### CAR assembly and constriction in cells of different geometry

To visualize actin cables, which are components of the CAR, we used a fluorescent mNeonGreen-tropomyosin probe, mNG-Tpm1 (Wirshing et al., 2025). A histone probe (H2B-mCherry) in the same strain revealed the timing of mitosis (Petrucco et al., 2024). In budding cells, CARs began to assemble following anaphase, 7 ± 0.8 min (mean ± sd, n = 200 cells) after mitotic entry as scored by chromosome condensation (Petrucco et al., 2024). In cells with multiple buds, all CARs constricted simultaneously (Fig. 2 and Video 1). Large multinucleated cells were uncommon in our culture conditions, but when cytokinesis was detected in such cells it was associated with assembly of very large CARs that constricted during septation (Fig. 2 and Video 1). In hyphae, mitosis often proceeded without cytokinesis, but on those occasions where cytokinesis did occur, CARs began to assemble 15 ± 5 min (mean ± sd, n = 203 septa) after mitotic entry and then constricted (Fig. 2 and Video 2). In addition, hyphal segments could form multiple buds, and the bud necks formed and constricted CARs simultaneously in a similar manner to multibudding cells (Video 2, left). Unlike the situation at bud necks, septa forming in the same hyphal segment were not always synchronous (Video 2, see upper right). Medial-plane kymographs indicated that CAR constriction proceeded at a constant rate, which was faster in hyphae than at bud necks (Fig. 2C: note that kymograph axes are equal for bud necks and hyphae). Although in *S. cerevisiae* cytokinesis can be accomplished in the absence of a CAR (Bi et al., 1998; Lord et al., 2005), all cytokinesis events that we observed in *A. pullulans* were associated with CAR assembly and constriction, regardless of the size or morphology of the cells.

**Figure 2:**
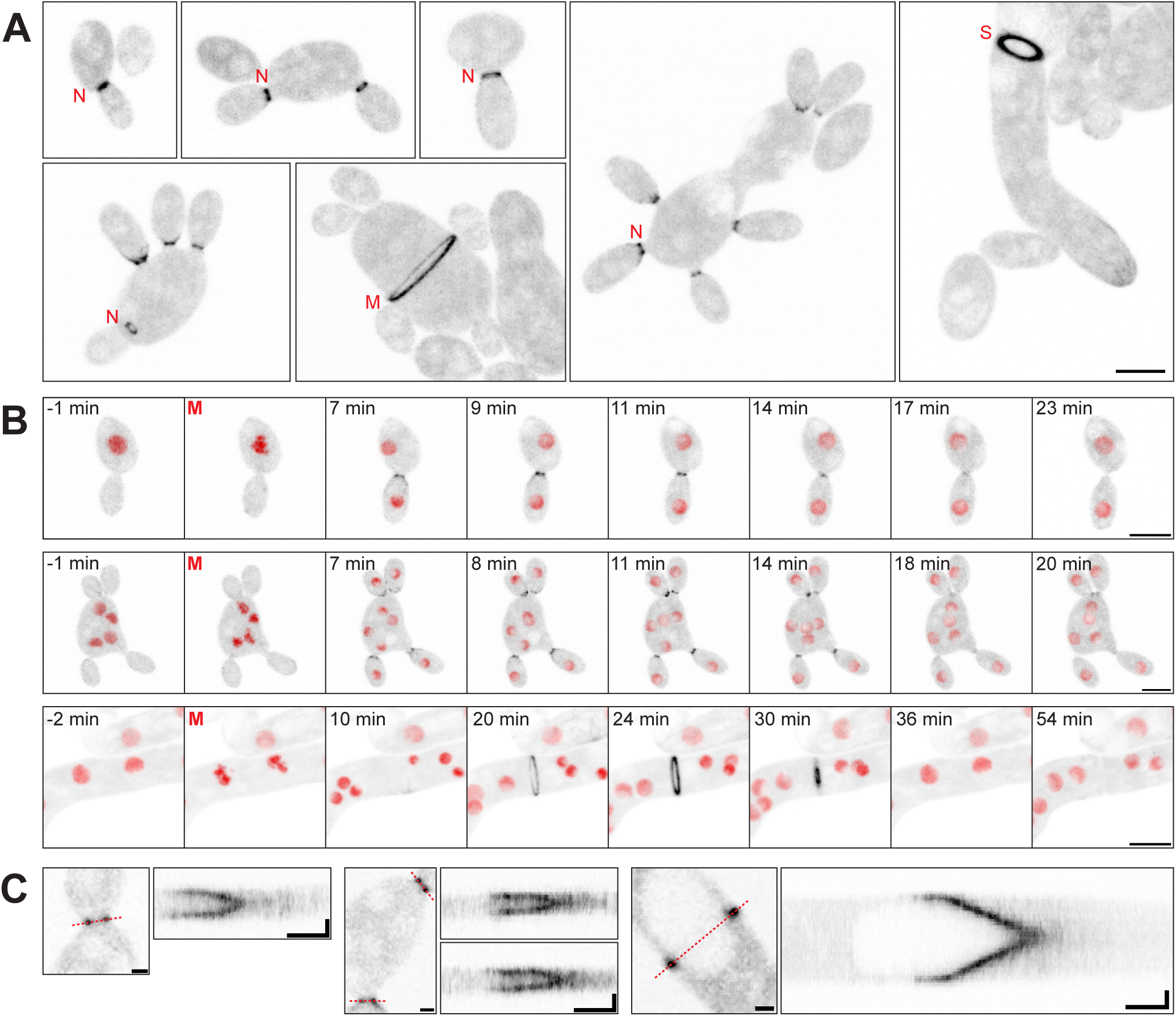
CAR assembly and constriction in *A. pullulans*. (A) CARs visualized with an actin-binding probe (mNG-Tpm1) in *A. pullulans* (strains DLY26075, DLY26077). Maximum projection inverted confocal images illustrating actin rings of different sizes at bud necks (N), hyphal septa (S), and medial locations in large multinucleate cells (M). Scale bar, 5 μm. (B) Time-lapse images of CAR formation and constriction in single-budded (top), multibudded (middle), and hyphal (bottom) cells imaged as in (A). The time of mitotic entry (chromosome condensation, visualized with H2B-mCherry probe in red) is designated as 0 min (red M). Scale bar, 5 μm. (C) Single-plane confocal images and associated kymographs illustrating CAR constriction at bud necks (left) and hyphal septa (right). The dashed red lines indicate the lines used to generate the associated kymograph. Scale bar, 1 μm. Kymograph axes 1 μm (vertical) and 5 min (horizontal).

### Septin localization in yeast

A previous study identified homologs of the four core septins, Cdc3, Cdc10, Cdc11, and Cdc12 in *A. pullulans* (Petrucco et al., 2024). A more extensive search identified a total of six septins in the *A. pullulans* genome, with single clear homologs of each core septin and two members of the filamentous-fungus-specific septin class defined by *A. nidulans* AspE (Pan et al., 2007)(Fig. 3A). In this study, we focus on the core septins. Heterologously expressed septins tagged with tdTomato could be seen at bud necks, although the signals were faint (Petrucco et al., 2024). In testing strategies for improved detection of septins, we found that heterologous expression could produce phenotypes suggestive of septation errors. However, C-terminal tagging of endogenous septin genes with mNG did not affect cell growth or morphology at 24°C, and only rarely produced errors in septation (Fig. S1). Using these strains, we were able to image septin localization in yeast (Fig. 3 and Video 3) and hyphae (Fig. 4 and Video 4) of *A. pullulans*.

**Figure 3:**
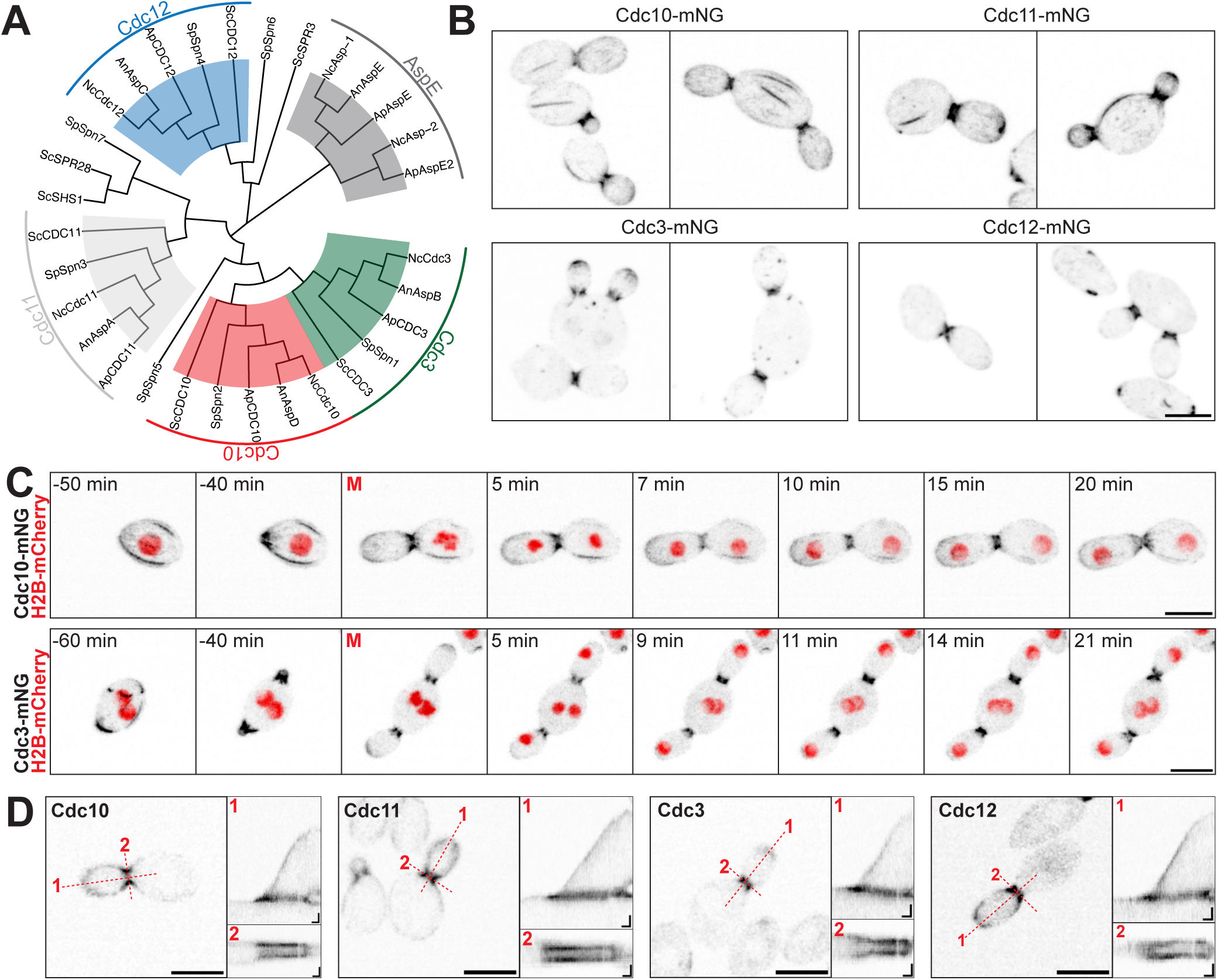
Septin localization in *A. pullulans* budding cells. (A) Evolutionary relationship between septins in *A. pullulans* (Ap) and those in the model ascomycetes *S. cerevisiae* (Sc), *S. pombe* (Sp), *A. nidulans* (An), and *N. crassa* (Nc). (B) Maximum projection inverted confocal images of budding cells expressing fluorescent septins (Cdc10-mNG, DLY26846; Cdc11-mNG, DLY26848; Cdc3-mNG, DLY26842; Cdc12-mNG, DLY26802). Scale bar, 5 μm. (C) Time-lapse maximum intensity projection images of cells expressing either Cdc10-mNG or Cdc3-mNG (black) and H2B-mCherry (red)(strains as in B). The time of mitotic entry is designated as 0 min (red M). Scale bar, 5 μm. (D) Single-plane confocal images and associated kymographs for each septin (strains as in B). The dashed red lines indicate the lines used to generate the associated kymograph. Scale bar, 5 μm. Kymograph axes 1 μm (vertical) and 5 min (horizontal).

**Figure 4:**
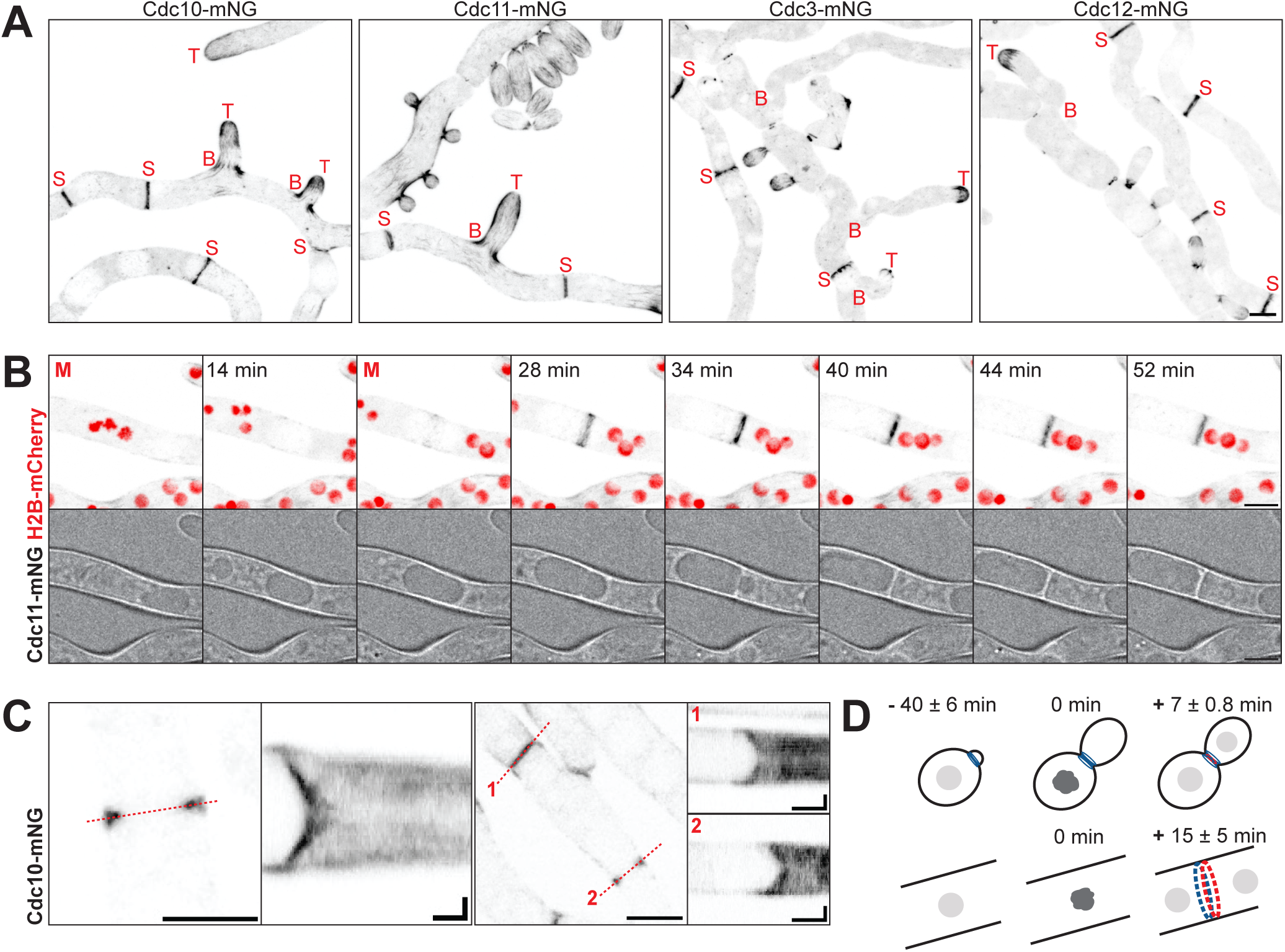
Septin localization in *A. pullulans* hyphae. (A) Maximum projection images of septins in hyphae (strains as in Fig. 3). Hyphal tips (T), septa (S), and branch sites (B) are indicated. Scale bar, 5 μm. (B) Time-lapse maximum intensity projection (top) and bright-field (bottom) images of a hypha expressing Cdc11-mNG (black) and H2B-mCherry (red)(DLY26848). The time of mitotic entry is designated as 0 min (red M). Scale bar, 5 μm. (C) Single-plane confocal images of septation sites and associated kymographs illustrating Cdc10-mNG dynamics during septation (DLY26846). The dashed red lines indicate the lines used to generate the associated kymograph. Scale bar, 5 μm. Kymograph axes 1 μm vertical, 5 min horizontal. (D) Schematic showing the average and standard deviation of the timing of actin and septin accumulation at the bud neck (n = 200 cells) or hyphal septum (septin n = 207 septa, actin n = 203 septa) relative to the timing of mitosis (0 min).

In yeast-form cells, all septins accumulated strongly at sites of bud emergence. A collar of septins stayed at each mother-bud neck throughout bud growth, while a more diffuse population of septins was present at the growing bud tip (Fig. 3B,C). At the time of mitosis, septin signals disappeared from bud tips and intensified briefly at bud necks (Fig. 3D and Video 3). After cytokinesis, septin remnants remained at the previous bud site for a variable period that could extend until after bud emergence in the next cell cycle. These findings indicate that as in *S. cerevisiae*, septins in *A. pullulans* are well placed to mark sites of future septation.

In addition to the localization pattern described above, which was common to all four core septins, the Cdc10-mNG and Cdc11-mNG probes often displayed cortical bars aligned with the mother-bud axis (Fig. 3B) that were rarely seen with Cdc3-mNG or Cdc12-mNG probes. The preferential decoration of cortical bars by Cdc10-mNG and Cdc11-mNG raised the possibility that there are structures that only incorporate a subset of septins. However, it was also possible that all structures contain full septin octamers, as they do in *S. cerevisiae*, but that tagging of particular septins affects the prevalence of the cortical bars (Gregory et al., 2025). In principle, these possibilities can be distinguished by tagging two septins with differing localization patterns in the same cell. For example, if cortical bars contain Cdc10 but not Cdc3, then one should detect bars only with the Cdc10 probe in double-tagged cells.

The localization pattern of each septin was unaffected by tagging with mScarlet instead of mNG (Fig. S2A). When different septins were tagged in the same cell, cortical bars always contained both of the tagged septins (Fig. S2B). These findings suggest that cortical bars are composed of full septin octamers. However, the prevalence of cortical bars was much lower in cells containing a tagged Cdc3 or Cdc12 probe than those containing only Cdc10 and Cdc11 probes (Fig. S2B). Thus, the prevalence of bars is affected by septin tagging.

Cells containing Cdc3-mNG or Cdc12-mNG probes displayed small rings and/or puncta that appeared to be released from the cytokinesis site to diffuse around the cell after septation (Video 3). Such puncta were rarely observed with Cdc10-mNG and Cdc11-mNG probes, and when cells contained two-color probe combinations, we frequently observed puncta that only contained a single (Cdc3 or Cdc12) probe (Fig. S2B). These puncta may therefore represent aggregates rather than intact septin structures.

### Septin localization in hyphae

In hyphae, all septins accumulated at growing hyphal tips and septation sites (Fig. 4A and Video 4). All septins also accumulated at hyphal branch sites during branch formation, but Cdc3-mNG and Cdc12-mNG dissipated quickly from those sites while Cdc10-mNG and Cdc11-mNG remained at the base of the branch for much longer (Video 4). As in budding cells (above), Cdc10 and Cdc11 also decorated filaments and bars aligned with the hypha axis at the hyphal cortex, while these were less evident with Cdc3-mNG and Cdc12-mNG probes (Fig. 4A and Video 4). The documented curvature preference of septins in other species (Bridges et al., 2016) was recapitulated in *A. pullulans*, particularly when imaging Cdc10-mNG and Cdc11-mNG. This was seen most clearly when hyphae curved and septins accumulated in the inner “elbow” (Video 4), although septin accumulation at such sites was transient.

During septation, septins localized to the ingressing furrow as well as the ingressed membrane (Fig. 4B,C). Septa forming in the same hyphal segment did not always constrict synchronously (Fig. 4C). After septum formation, septins disassembled with variable timing. Small septin rings and puncta dissociated from the septation site and sometimes persisted for some time as mobile structures in the nearby hyphal shaft, particularly with the Cdc12-mNG probe (Video 4).

The timing of septin accumulation at septation sites was similar to that of actin, with both septin and actin rings assembling 15 ± 5 min (mean ± sd, n = 167-207 septa) after mitotic entry (Fig. 4D). Thus, whereas septins are present and marking the bud neck for about an hour before cytokinesis in yeast-form cells, they appear to arrive at hyphal septation sites at the same time as the CAR just before cytokinesis. These findings are consistent with a role for septins in CAR placement at bud necks but perhaps not hyphal septation sites.

### Stochastic cytokinesis failure in cells lacking individual septins

To address the role(s) of septins in cytokinesis, we deleted each of the four core septin genes. Septin deletions were well tolerated at 24°C, although colony growth was temperature sensitive (Fig. S1). Because of the septin localization data discussed above, we initially focused on cytokinesis at the mother-bud necks of yeast-form cells. To our surprise, most mutant cells successfully assembled and constricted CARs at mother-bud necks (Fig. 5 and Video 5). However, a subset of necks did not assemble CARs and failed to undergo cytokinesis, yielding chains of connected cells (Fig. 5 and Video 5). Individual multibudded cells assembled CARs at some but not other bud necks (indicated by arrowheads vs asterisks in Fig. 5A,B), showing that the same cell could display both successful and unsuccessful cytokinesis sites. There was no difference in neck diameter between necks that did or did not assemble a CAR (Fig. S3A,B), and in some cases a neck that failed to assemble a CAR in one cell cycle succeeded in assembling a CAR in the next cell cycle (Fig. S3C). These observations suggest that cytokinesis failure may not be driven by a pre-existing defect at particular necks. Instead, septin mutants display a sporadic, apparently stochastic defect in the ability to assemble a CAR at mother-bud necks.

**Figure 5:**
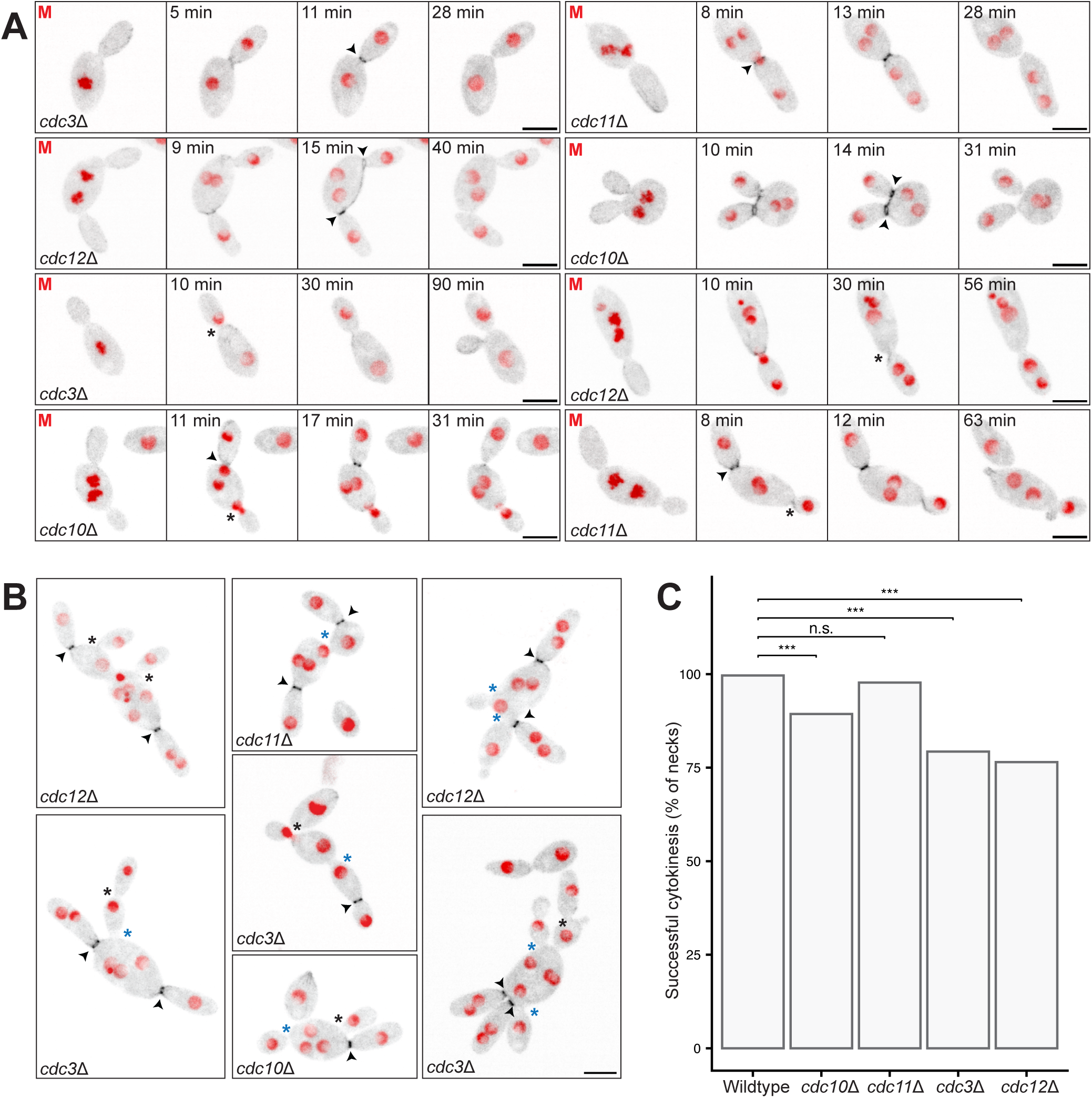
Cytokinesis in septin mutants. (A) Time-lapse maximum intensity projection confocal images of actin cables (mNG-Tpm1, black) and nuclei (H2B-mCherry, red) in the indicated septin mutants undergoing cytokinesis (two independent transformant strains imaged for each deletion: *cdc3*Δ, DLY26176 & DLY26177; *cdc10*Δ, DLY26179 & DLY26180; *cdc11*Δ, DLY26182 & DLY26183; *cdc12*Δ, DLY26185 & DLY26186). Arrowheads: bud necks that underwent successful cytokinesis. Asterisks: bud necks that failed to undergo cytokinesis. The time of mitotic entry is designated as 0 min (red M). Scale bar, 5 μm. (B) Examples of septin mutant chaining following failed cytokinesis. Arrowheads and asterisks as in (A). Blue asterisks indicate bud necks that failed to undergo cytokinesis in the previous cell cycle. Scale bar, 5 μm. (C) Cytokinesis success rate as a percent of bud necks (n = 270, 253, 265, 284, 285 necks for WT, *cdc10*Δ, *cdc11*Δ, *cdc3*Δ, and *cdc12*Δ mutant strains). Statistical significance calculated by Chi-squared test (n.s., p > 0.05; ***, p ≤ 0.005).

Septation defects were most severe in *cdc3*Δ and *cdc12*Δ mutants, where 25% of necks failed to assemble CARs, and mildest in *cdc11*Δ mutants (Fig. 5C). The difference in severity suggests that remaining septins may retain partial function in *cdc10*Δ and *cdc11*Δ mutants, which is supported by the data presented below.

In addition to septation defects, a minority of septin mutant cells either stopped growth of a small bud or continued growth of a larger bud to make unusually elongated buds (most severe for *cdc3*Δ and *cdc12*Δ mutants: Fig. S4A-C). These phenotypes suggest a potential role for septins in controlling bud growth and morphology. We also observed sporadic defects in the behavior of nuclei. In wild-type cells, nuclei remain in the mother during bud growth and only transit the necks into buds during mitosis (Petrucco et al., 2025). However, in septin mutants, nuclei sometimes entered the buds during interphase (particularly for cells with large elongated buds) (Fig. S4C,D). Moreover, during mitosis some nuclei appeared to become stuck within the neck, and occasionally nuclei returned from the neck or the bud to the mother, leaving an anucleate bud (Fig. S4C,D). These nuclear defects were most pronounced in *cdc10*Δ mutants, and they suggest that septins play roles in nuclear movement as well as cytokinesis.

### Septin localization in cells lacking other septins

The differences in the penetrance of various septin mutant phenotypes suggested that when one septin is absent, the remaining septins may retain the ability to associate and perform some functions. Consistent with that idea, previous studies in *S. cerevisiae* showed that cells lacking Cdc10 or Cdc11 could still form partially functional septin hexamers capable of polymerizing to form filaments (McMurray et al., 2011). To investigate whether the remaining septins might localize to the mother-bud neck to enable cytokinesis in *A. pullulans* septin mutants, we imaged each septin in strains deleted for every other septin (Fig. S5). Cells lacking Cdc10 or Cdc11 were able to recruit reduced amounts of each remaining septin to some, though not all, bud necks (Fig. S5). However, cells lacking Cdc3 or Cdc12 did not recruit detectable amounts of any other septin to bud necks (Fig. S5). Instead, the other septins were distributed diffusely throughout the cytoplasm (Cdc3, Cdc10) or in multiple small diffusing puncta (Cdc11, Cdc12).

To assess whether cells might transiently recruit septins to the bud neck, we performed a time-lapse analysis of Cdc10-mNG (Fig. 6A) and Cdc12-mNG (Fig. 6B) localization in the other septin mutants. Both septins could localize to the neck in *cdc11*Δ mutants, albeit in a more irregular, faint, and transient manner than in wild-type cells. Similarly, Cdc12 could localize to the neck in some *cdc10*Δ mutant cells. However, no neck localization was detected for either probe in *cdc3*Δ mutants, and no neck localization of Cdc10 was detected in *cdc12*Δ mutants (Fig. 6). We conclude that septin subcomplexes can form and assemble at bud necks in the absence of Cdc10 or Cdc11, but not in the absence of Cdc3 or Cdc12. Thus, CAR assembly and cytokinesis can succeed in mutants (*cdc3*Δ, *cdc12*Δ) that have no detectable septin structures at the neck.

**Figure 6:**
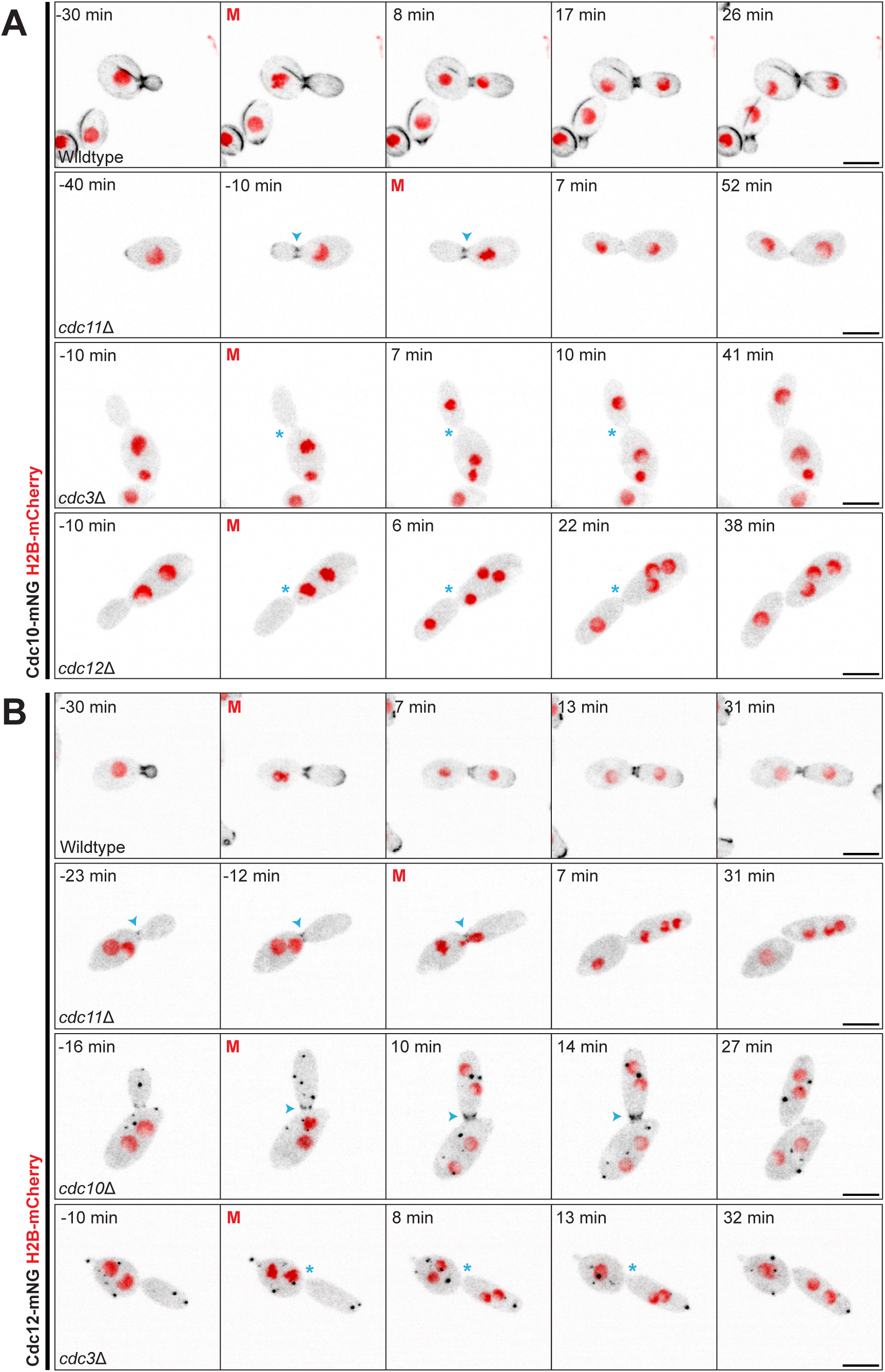
Septin localization in the absence of other septins. Time-lapse maximum intensity projection confocal images of cells with the indicated septin deletions. (A) Cdc10-mNG (black) and H2B-mCherry (red). No localized Cdc10 was detected at the bud necks of *cdc3*Δ (DLY27070) or *cdc12*Δ (DLY27076) mutants (blue asterisks), but Cdc10 was often faintly present at bud necks of *cdc11*Δ mutants (DLY27073)(blue arrowheads). (B) Cdc12-mNG (black) and H2B-mCherry (red). Cdc12 was predominantly detected in small puncta in the cytoplasm. No Cdc12 was detected at bud necks of *cdc3*Δ (DLY27061) mutants (blue asterisks), but Cdc12 was sometimes faintly present at bud necks of *cdc10*Δ (DLY27065) and *cdc11*Δ (DLY27067) mutants (blue arrowheads). The time of mitotic entry is designated as 0 min (red M). Scale bar, 5 μm.

### Core septins are dispensable for cytokinesis

Our experiments thus far suggest that while septins can improve the efficiency with which CARs assemble at bud necks, organized septin scaffolds are dispensable for CAR assembly and function at a majority of bud necks. However, there remained the possibility that delocalized septins, or septin structures too faint or transitory to detect in our imaging conditions, might contribute to cytokinesis. To definitively investigate this possibility, we generated a strain lacking all four core septin genes. This strain exhibited bud elongation, cytokinesis, and nuclear behavior defects with penetrance comparable to that of the most severe single mutant (Fig. 7 and Fig. S4). Nevertheless, a majority (75%) of bud necks assembled correctly-placed CARs and underwent successful cytokinesis (Fig. 7A,C). We conclude that *A. pullulans* yeast cells must have a mechanism to identify the bud neck as a site for CAR placement even in the absence of septins.

**Figure 7:**
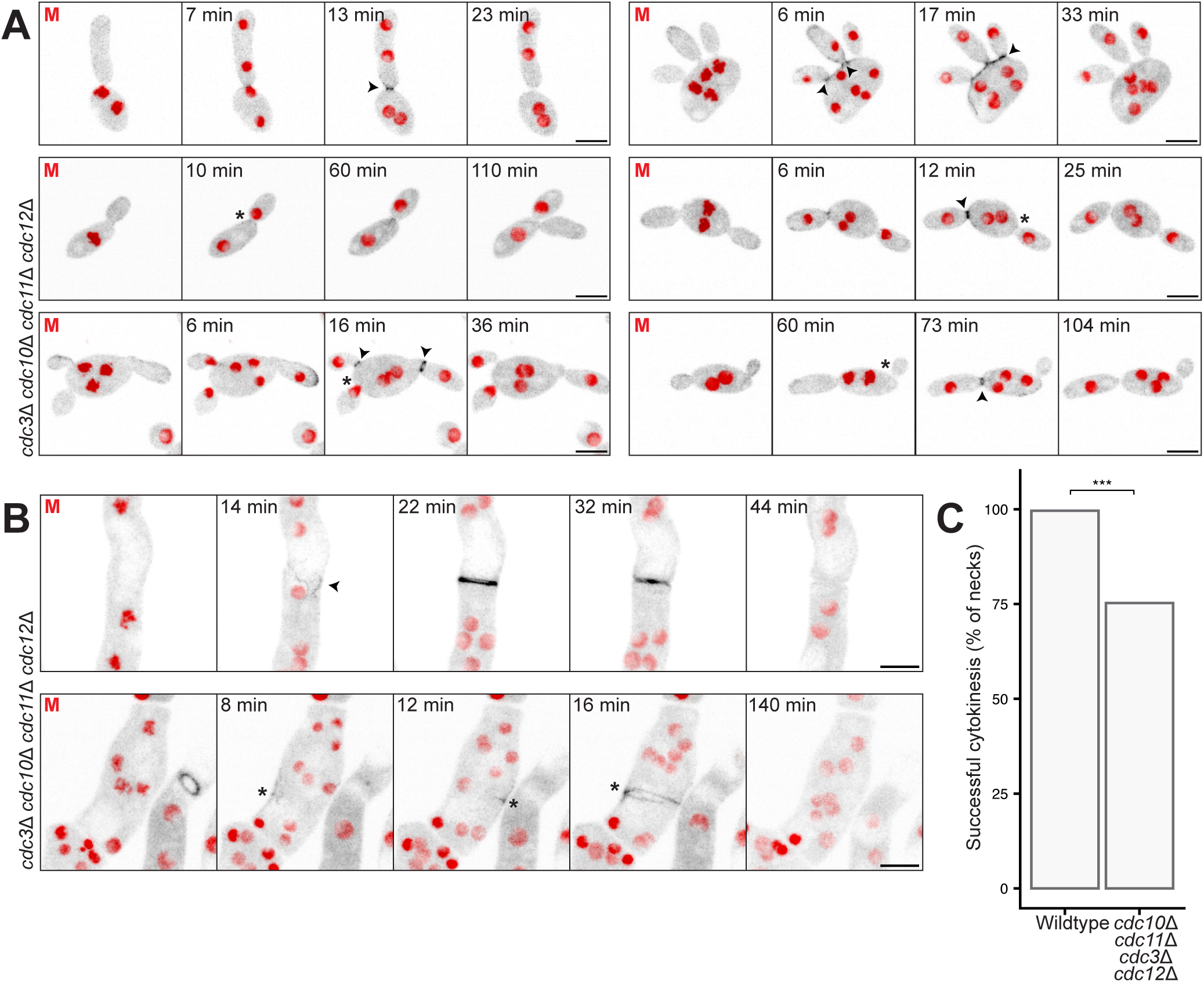
Cytokinesis in cells lacking all core septins. Time-lapse maximum intensity projection confocal images of actin cables (mNG-Tpm1, black) and nuclei (H2B-mCherry, red) in strains with all four core septin genes deleted (two independent transformant strains imaged: DLY27187, DLY27188). (A) budding cells. (B) hyphae. Arrowheads: bud necks and hyphal septa that underwent successful cytokinesis. Asterisks: bud necks and hyphae that failed to undergo cytokinesis. The time of mitotic entry is designated as 0 min (red M). Scale bar, 5 μm. (C) Cytokinesis success rate as a percent of bud necks (n = 270 necks for WT, 283 necks for mutant). Statistical significance calculated by Chi-squared test (***, p ≤ 0.005).

In hyphae, the mutant lacking all four core septins was still able to form and constrict CARs, leading to successful septation (Fig. 7B, upper panels). However, there were also instances in which mutant cells appeared to initiate CAR formation but failed to complete or constrict the CARs (Fig. 7B, lower panels). Thus, septins likely play ancillary roles in septation of hyphae as well as yeast.

### Cytokinesis in the combined absence of septins and mitotic nuclear cues

Previous studies showed that cytokinesis could occur at the bud neck even in cases where no nucleus was inherited by the bud, suggesting that spatial cues from mitotic nuclei are not required for successful cytokinesis (Petrucco et al., 2025). However, it seemed possible that septins and mitotic nuclei (or spindles) might provide partially redundant CAR localization cues. To assess whether that might be the case, we asked whether migration of a mitotic nucleus through the neck was required to promote CAR assembly in cells that lacked Cdc12 (where no septins are visible at the neck).

We used the microtubule depolymerizing drug benomyl to block chromosome segregation and nuclear movement, keeping nuclei in the mother cell. As in other fungi, benomyl treatment arrested cells in mitosis with clearly condensed chromosomes, presumably via the spindle assembly checkpoint (Lew and Burke, 2003) (Fig. 8A). To allow mitotic exit and cytokinesis in benomyl-treated cells, we deleted the checkpoint component *MAD2* (Lew and Burke, 2003). *mad2*Δ mutants were viable and proliferated well in the absence of benomyl, but failed to arrest upon microtubule depolymerization (Fig. 8A). Instead, chromosomes condensed and then decondensed without forming a spindle (Fig. 8A). *cdc12*Δ *mad2*Δ double mutants treated with benomyl did not segregate nuclei through the bud neck, but most necks nevertheless assembled CARs and successfully completed cytokinesis, yielding anucleate buds (Fig. 8B,C). We conclude that CAR assembly and constriction at the mother-bud neck can proceed in the absence of septins and without passage of mitotic nuclei through the neck, suggesting that *A. pullulans* cells possess a novel mechanism to specify the neck as a cleavage site.

**Figure 8:**
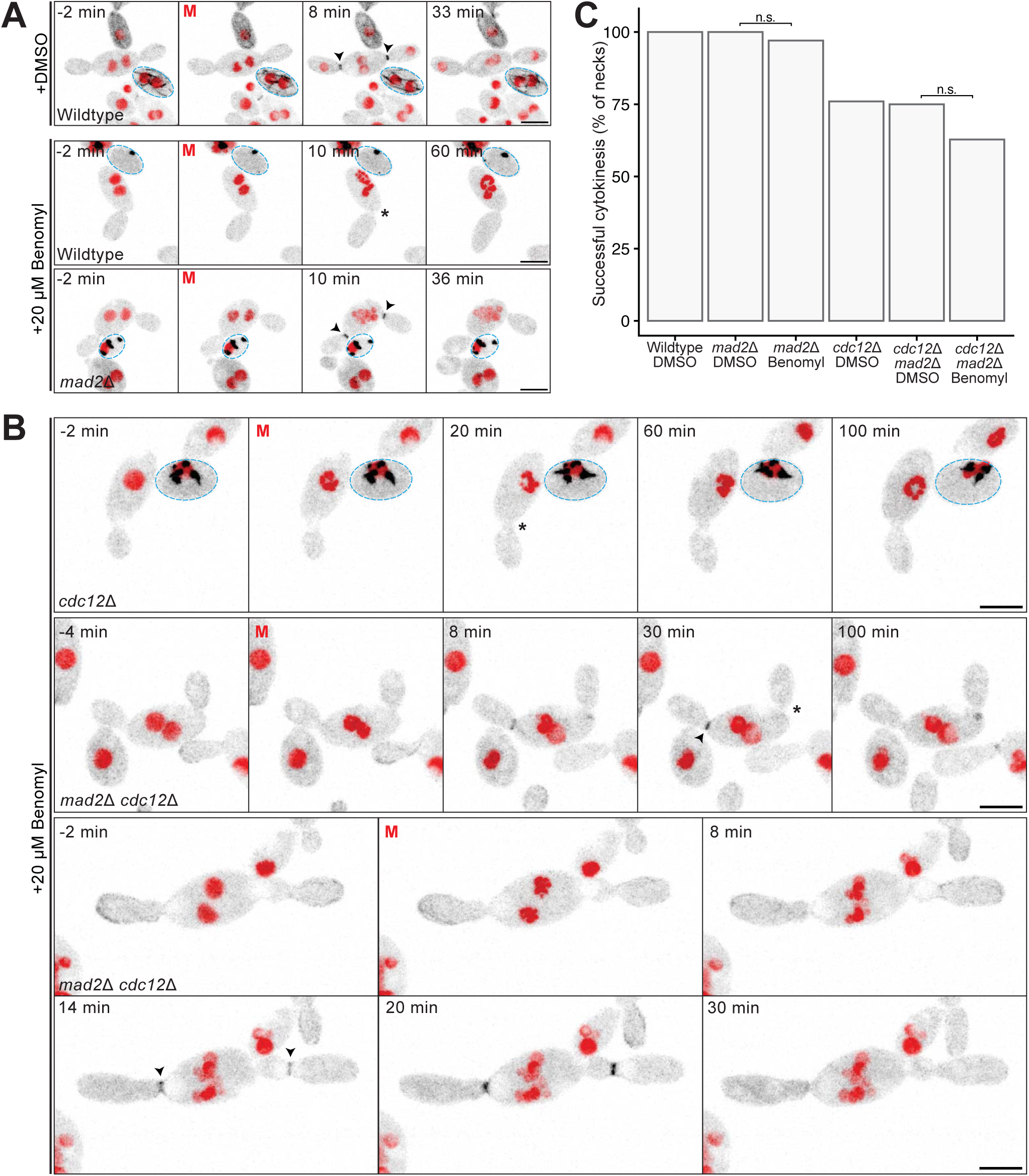
Cytokinesis in cells lacking polymerized septins and microtubules. (A) Time-lapse maximum intensity projection confocal images of actin cables (mNG-Tpm1, black) and nuclei (H2B-mCherry, red) in Wildtype (DLY26075) or *mad2*Δ (DLY27434) strains treated with DMSO (control) or 20 μM Benomyl to depolymerize microtubules. The effectiveness of the benomyl treatment was assessed by spiking in cells that express 3xGFP-Tub2 (black) and H2B-mCherry (red): these cells are outlined in blue (DLY24568). Scale bar, 5 μm. (B) Similar images for *cdc12*Δ (DLY26185) and *cdc12*Δ *mad2*Δ (DLY27437) cells treated with 20 μM Benomyl. With an intact spindle assembly checkpoint (MAD2), cells arrest with condensed chromosomes and do not assemble CARs (top row, asterisk). In contrast, checkpoint mutants (*mad2*Δ) condense and then decondense their chromosomes without assembling a spindle or delivering nuclei to buds. Nevertheless, these cells often complete cytokinesis (arrowheads). The time of mitotic entry is designated as 0 min (red M). Scale bar, 5 μm. (C) Cytokinesis success rate as a percent of bud necks (n = 40, 46, 34, 50, 48, and 43 necks for WT DMSO, WT *mad2*Δ DMSO, WT *mad2*Δ benomyl, *cdc12*Δ DMSO, *cdc12*Δ *mad2*Δ DMSO, *cdc12*Δ *mad2*Δ benomyl). Statistical significance calculated by Chi-squared test (n.s., p > 0.05).

## DISCUSSION

Mechanisms of CAR placement and constriction can vary widely between species. For example, nuclear cues direct the placement of a CAR in the medial region of *S. pombe*, which assembles and constricts in a manner that requires myosin motor activity (Kitayama et al., 1997; Bezanilla et al., 1997; Balasubramanian et al., 1998; Bähler et al., 1998; Wu et al., 2006; Almonacid et al., 2009; Saha and Pollard, 2012). In contrast, septin scaffolds direct the placement of a CAR at the much narrower bud neck in *S. cerevisiae*, which can assemble and constrict without myosin motor activity (Longtine et al., 1996; Bi et al., 1998; Lord et al., 2005; Fang et al., 2010). It is unclear to what degree these differences reflect the differing needs of larger versus smaller CARs, or the different constraints imposed by tubular versus locally constricted cell geometry, or simply the divergent evolutionary histories of the two yeasts. We show that *A. pullulans* cells of widely different sizes and shapes all use CARs during cytokinesis, establishing *A. pullulans* as a model for addressing how CAR placement, assembly, and constriction mechanisms might vary as a function of cell size or shape. As a first step towards answering these questions, we investigated the roles of septins in *A. pullulans*.

### Septin localization in *A. pullulans*

The *A. pullulans* genome contains a single homolog of each of the four core septin genes characterized in *S. cerevisiae*: Cdc3, Cdc10, Cdc11, and Cdc12. We did not detect a homolog of the nonessential *S. cerevisiae* septin Shs1/Sep7, but there were two homologs of the *A. nidulans* septin AspE, a noncore septin found only in filamentous fungi (Pan et al., 2007) (Fig. 3A). Core septins in *S. cerevisiae* assemble into rod-shaped non-polar octamers (Cdc11-Cdc12-Cdc3-Cdc10-Cdc10-Cdc3-Cdc12-Cdc11) that can associate with the plasma membrane and polymerize to form various structures (Frazier et al., 1998; Bertin et al., 2008; Bridges et al., 2016; Ong et al., 2014). Septins have an intrinsic preference for associating with membranes that have shallow micron-scale curvature, such as those found at mother-bud necks and hyphal branch sites (Bridges et al., 2016). This preference has been traced to amphipathic helices found at the C-termini of some septins (Cannon et al., 2019; Woods and Gladfelter, 2021). Consistent with a similar behavior for *A. pullulans* septins, all of the core septins localized to emerging buds and mother-bud necks, as well as hyphal branches. When hyphae curved, septins frequently accumulated at the “inner elbow” of the curve, which has similar curvature to that of bud necks and hyphal branches.

Additionally*, A. pullulans* septins accumulated at growing bud tips and hyphal tips. These sites are associated with polarity regulators (Crocker et al., 2025), which recruit septins in *S. cerevisiae* (Pringle et al., 1995; Gladfelter et al., 2002; Caviston et al., 2003). Thus, it seems likely that *A. pullulans* polarity factors first recruit septins to growth sites and then leave them behind at sites with the favored membrane curvature (bud necks and branch sites).

In *A. pullulans* hyphal cells or large multi-nucleate cells, septins were recruited to the site of septation concomitant with division. Unlike at bud necks, these sites lack positive membrane curvature. Thus, other signals likely direct septin localization. In the tube-shaped *S. pombe* cells, mitotic factors including Mid2 (anillin homologue) recruit septins to a ring at the site of division (An et al., 2004). Similarly, in hyphal *A. nidulans* cells septin recruitment to septation sites requires CAR proteins SepA/formin, SepG/IQGAP, and SepH/Hippo-like kinase (Westfall and Momany, 2002). It will be interesting to determine if *A. pullulans* uses similar or distinct strategies to direct the positioning of septins to septation sites.

### Composition of septin structures in *A. pullulans*

While all four core septins were present at growing tips, bud necks, and hyphal branch sites, other structures were visualized to different extents using different septin probes. In particular, cortical septin filaments and bars were more prevalent in cells carrying Cdc10 and Cdc11 probes, while small rings and puncta were most prevalent in cells with the Cdc12 probe. The specific fluorophore (mNG or mScarlet) did not appear to affect localization. Our analysis of cells bearing pairs of probes indicated that the cortical filaments and bars contained combinations of all four core septins, while the cytoplasmic puncta often did not. We speculate that cortical filaments and bars represent native structures made from standard septin octamer subunits, while cytoplasmic puncta may represent unnatural aggregates of tagged septins. In addition, the prevalence of the cortical filaments and bars was affected by tagging of specific septins, so the true prevalence of such structures remains unclear. The potential for septin tagging to affect localization and function was recently explored in some depth in *S. pombe* (Gregory et al., 2025), and our findings are consistent with their conclusions. Overall, our results are consistent with the idea that as in model yeasts, physiological septin structures in *A. pullulans* are formed from protomers that contain each of the four core septins.

The phenotypes of individual septin mutants ranged in severity, with *cdc3*Δ and *cdc12*Δ being most severe and *cdc11*Δ least severe. *cdc10*Δ and *cdc11*Δ mutants retained the ability to localize other septins to the mother-bud neck, albeit to a reduced degree. These findings are most easily interpreted as indicating that protomers composed of Cdc10-Cdc3-Cdc12 (in *cdc11*Δ mutants) or Cdc3-Cdc12-Cdc11 (in *cdc10*Δ mutants) retain some ability to polymerize, and hence provide partial function, consistent with previous work (McMurray et al., 2011). In contrast, *cdc3*Δ and *cdc12*Δ mutants did not detectably localize other septins to their normal locations, and they were phenotypically similar to quadruple *cdc3*Δ *cdc10*Δ *cdc11*Δ *cdc12*Δ mutants. Thus, with the caveat discussed below, *cdc3*Δ and *cdc12*Δ mutants appear to lack all core septin function.

Interestingly, septin mutants also had defects in nuclear inheritance during mitosis. In particular, some mothers failed to deliver a nucleus through the neck into a bud, while in other cases a mitotic nucleus appeared to become stuck during transit through the neck. These phenotypes were most prevalent in *cdc10*Δ mutants, perhaps suggesting that Cdc10 has a specific role in nuclear migration, that does not require its association with the other septins. Interestingly, one of the isoforms of mammalian Septin 9 contains a microtubule-binding domain at its N-terminus (Kuzmić et al., 2022). However, *A. pullulans* Cdc10 lacks this domain, and the significance of the nuclear migration phenotype remains unclear.

### Septins and cytokinesis

The major defect in septin mutants was the failure of CAR assembly at a subset (about 25%) of mother-bud necks. This supports the hypothesis that as in *S. cerevisiae*, septins in *A. pullulans* act as a scaffold that locally concentrates CAR components to promote cytokinesis. Interestingly, there was an additional burst of septin recruitment to the bud neck at the time of cytokinesis in *A. pullulans*, whereas in *S. cerevisiae* septins are partially released from the neck at that time (Ong et al., 2014). This may indicate divergent roles of septins in the two yeasts. In *S. cerevisiae*, septins sequentially recruit different CAR components during bud growth, beginning with myosin II and later IQGAP before the final recruitment of actin cables upon mitotic exit (Bhavsar-Jog and Bi, 2017; Howell and Lew, 2012). Here we only examined actin cables, which assembled following mitotic exit in *A. pullulans*. It will be interesting to determine whether CAR assembly follows a similar or distinct temporal order in *S. cerevisiae* and *A. pullulans*.

In *A. pullulans* hyphae, septins were only recruited to septation sites at the time of cytokinesis, coincident with CAR assembly. Thus, as in *A. nidulans* (Seiler and Justa-Schuch, 2010), septins are unlikely to direct CAR placement in hyphae. Septin mutants could still assemble and constrict CARs in hyphae, although possibly with reduced efficiency. Future work will explore the roles of septins in hyphae of *A. pullulans*.

### Targeting CAR assembly

The most surprising finding from this study was that CARs still assembled and constricted at 75% of bud necks in septin mutants. This reveals the unexpected existence of a second pathway to target CAR assembly to the neck. Using microtubule poisons, we found that this second pathway was able to function even in the combined absence of septins and microtubules, indicating that it does not rely on nuclear passage through the bud neck. How cells are able to target CAR assembly (albeit less efficiently) in such cells is unknown. We speculate that there may be another, septin-independent way to recognize the unusual membrane curvature present at the bud neck (Kang et al., 2016) in order to target CAR assembly at that site.

Septin-independent cytokinesis pathways may also be present in other budding yeasts. Notably, mutants deleted for individual septin genes were viable (although temperature sensitive) in *Ustilago maydis* (Alvarez-Tabarés and Pérez-Martín, 2010) and *Cryptococcus neoformans* (Kozubowski and Heitman, 2010). This suggests that as in *A. pullulans*, cytokinesis can proceed without septins in these basidiomycete yeasts. However, unlike in *A. pullulans*, some septin double mutants were lethal in *U. maydis*, raising the possibility that viable mutants retained some septin function (Alvarez-Tabarés and Pérez-Martín, 2010). It will be interesting to determine whether septin-independent CAR targeting mechanisms are conserved in other budding yeasts.

Beyond budding yeasts, there remain significant mysteries with regard to how cells target CAR assembly. In filamentous fungi, it is unclear how CAR locations in hyphae are determined. In animal cells it is generally accepted that the mitotic spindle instructs CAR location in most cases (Glotzer, 2017), but specialized cells like the pole cells in *Drosophila* embryos assemble CARs to allow budding off from the syncytial embryo (Cinalli and Lehmann, 2013), and the mechanisms instructing CAR locations in this case are mysterious. As a polymorphic fungus that assembles CARs in several different morphologies, *A. pullulans* provides an opportunity to uncover novel CAR targeting pathways that may be applicable in other contexts.

## METHODS

### *A. pullulans* strains

All *A. pullulans* strains used in this study are listed in Supplemental Table 1 and were constructed in the *EXF-150* strain background (Gostinčar et al., 2014). Strains were grown at 24°C in YPD (2% glucose, 2% peptone, 1% yeast extract) on plates solidified with 2% BD Bacto^TM^ agar (214050, VWR).

To express *A. pullulans* tropomyosin (Tpm1) tagged at its N-terminus with mNeonGreen, mNG-Tpm1 with its native promoter and terminator was amplified from DLB4897 (Wirshing et al., 2025) and inserted into DLB4690 to generate DLB4995, which contains *A. pullulans URA3* and 1kb flanking homology regions. PCR products were amplified using Phusion™ Hot Start Flex 2X Master Mix (M0536L, NEB) following manufacturer’s instructions. Fragments were assembled with NEBuilder HiFi DNA Assembly Master Mix (E2621L, New England Biolabs).

To express both histone and tropomyosin markers, mNG-Tpm1 with 1kb downstream *URA3* homology was amplified from DLB4995, and SpH2B-mCherry with *URA3* and 1kb upstream homology was amplified from DLB4847 (Wirshing et al., 2025). Primers were designed to introduce 66 bp overlap to allow for homologous recombination in the yeast. The PCR fragments were directly added to competent cells, generating DLY24148.

Tagging of endogenous Cdc3 (protein ID 275598), Cdc10 (protein ID 342729), Cdc11 (protein ID 354011), and Cdc12 (protein ID 344134) with mNG was performed as described (Colarusso et al., 2025) with mNG amplified from pAPInt-mNeonGreen-Nat (DLB4908, Addgene #236482). Septin and Mad2 (protein ID 184083) deletions were also constructed as described (Colarusso et al., 2025), by deleting the entire open reading frame and replacing it with a selectable marker. Genes were deleted using selection with nourseothricin, hygromycin, geneticin, and phleomycin (Ble). To use Ble selection, a pAPInt vector was constructed with the Ble resistance gene, *Sh ble*, amplified from plasmid pRS40B-bleMX4 (Chee and Haase, 2012) and inserted into the pAPInt-GFP-Nat backbone (DLB4760, Addgene #236476). The appropriate concentration of phleomycin (P9564, Millipore) was determined by plating 10^7^ cells onto YPD plates supplemented with concentrations ranging from 0 to 100 µg/ml. Selection was performed on plates with 10 µg/ml phleomycin.

All *A. pullulans* strains were constructed using the chemical transformation method as previously described (Wirshing et al., 2024). Transformants expressing fluorescent probes were screened by microscopy and confirmed using colony PCR to check for correct integration of fluorescent tags or deleted genes.

To check the effect of tags or deletions on colony growth, a single colony was inoculated into 5 ml YPD (2% glucose) and grown for 2 days at 24°C. Serial 1:10 dilutions of cultures with sterile water were plated onto YPD plates with a pin-frogger. Plates were grown for 2 days at 24°C or 30°C and imaged on an Amersham Imager 680 (General Electric Company) with colorimetric epi settings.

### Live-cell imaging

To image budding cells, cells were grown overnight at 24°C in YPD (2% glucose) to a density of 1-5 x 10^6^ cells/ml, pelleted for 10 s at 9391 rcf, and resuspended to a density of 5-10 x 10^7^ cells/ml. 1-2 x 10^5^ cells were plated onto an 8-well glass-bottomed Ibidi chamber (80827, Ibidi) and covered with a pad of complete synthetic media (CSM: 6.71 g/L BD DifcoTM Yeast Nitrogen Base without Amino Acids, BD291940, FisherScientific, 0.79 g/L Complete Supplement Mixture, 1001-010, Sunrise Science Products, and 2% glucose) solidified with 1.5% agarose (97062-250, VWR).

To image hyphae, cells were grown overnight in liquid CSM without agitation at 24°C. 2-3 x 10^6^ cells were plated onto an 8-well glass-bottomed Ibidi chamber (80827, Ibidi), covered with a CSM pad solidified with 5% agarose, and grown overnight at 24°C. Hyphal networks were imaged the following morning.

To image larger multinucleate cells, cells were grown overnight at 24°C in YPD (2% glucose) to a density of 1 x 10^6^ cells/ml, pelleted at 10 s at 9391 rcf, and resuspended to a density of 5 x 10^7^ cells/ml. Cells were then left to settle for 30 min in a 1.5 ml epi tube. A pipette was used to carefully collect cells from the bottom of the tube, and 1-2 x 10^5^ cells were plated onto a CSM pad solidified with 1.5% agarose.

All imaging was conducted on a Nikon Ti2E inverted microscope with a CSU-W1 spinning-disk head (Yokogawa), CFI60 Plan Apochromat Lambda D 60x Oil Immersion Objective (NA 1.42; Nikon Instruments), and a Hamamatsu ORCA Quest qCMOS camera controlled by NIS-Elements software (Nikon Instruments). Cells were imaged at room temperature (20-22°C). Still images were acquired at 0.2 µm step intervals for a total of 75 Z-slices.

For time-lapse imaging of budding cells, stacks of 19 Z-slices at 0.4 or 0.5 µm step intervals were acquired at 1 min intervals for 2-3 h. For hyphae, stacks of 21 Z-slices at 0.6 µm step intervals were acquired at 2 min intervals for 2-3 h. For mNG-Tpm1 kymographs, 19 or 21 Z-stacks at 0.5 µm step intervals were acquired at 10 s intervals for a total of 45 min. For septin-mNG kymographs, 19 or 21 Z-stacks at 0.5 µm step intervals were acquired at 30 s intervals for 1 h.

Cytokinesis in cells lacking septins and mitotic nuclear cues were observed by treating cells with benomyl (20 µM final concentration). CSM pads solidified with 5% agarose were incubated in CSM with the final benomyl (20 mM stock in DMSO) concentration. Pads for control experiments were incubated with the same volume of DMSO. The pads were then dried briefly on a kimwipe before being placed on cells. 13 Z-stacks at 0.5 µm step intervals were acquired at 2 min intervals for 3 h.

Exposure times were 50 ms at 15-20% laser power for mNG-Tpm1 and 50 ms at 10-20% laser power for septin-mNG (excitation 488 nm). Exposure times were 50 ms at 20-30% laser power for septin-mScarlet and 50 ms at 8-10% laser power for histone-mCherry (excitation 561 nm).

### Image analysis

Images were median denoised using NIS Elements (Nikon Instruments). All image analysis was done in FIJI (Schindelin et al., 2012) unless indicated otherwise.

Scoring of the intervals between mitosis and CAR assembly at the bud neck or septum was done using time-lapse Z-stack images of cells expressing mNG-Tpm1 and histone-mCherry. Mitosis was scored as the time of histone condensation as in (Petrucco et al., 2025), and ring assembly was scored as the first frame where mNG-Tpm1 was visible at the neck or septum. Scoring for timing of bud emergence, mitosis, and septin localization was done using time-lapse Z-stack images of cells expressing septin-mNG and histone-mCherry. Bud emergence/septin arrival at the neck was scored by septin concentration to a bud site and confirmed by scoring a visible bud in the brightfield image.

Kymographs of CAR and septin dynamics over time were generated in FIJI using a single medial plane. Images were first processed with StackReg plugin in FIJI to correct for drift (Thevenaz et al., 1998).

To measure circularity of buds, confocal Z-stacks were median denoised and maximum intensity projected in NIS Elements (Nikon Instruments), then binarized in FIJI. Buds were selected at the time point after cytokinesis, and the circularity was measured using FIJI. To measure bud neck diameter, the full width at half maximum at the time of mitosis was measured using cytoplasmic Tpm1 signal.

### Phylogenetic tree

*A. pullulans* septins were identified using the basic local alignment search tool (BLAST) against *A. pullulans* EXF-150 through JGI MycoCosm. Septin protein sequences were aligned using Clustal Omega and visualized with R package ggtree (Yu et al., 2017).

### Statistical analysis

All statistical analysis was completed in R using the stats package.

## DATA AVAILABILITY

The data generated in this study are available from the corresponding authors upon reasonable request.

## ACKNOWLEDGEMENTS

We thank Amy Gladfelter (Duke University), Mohan Balasubramanian (Warwick University, UK), Masayuki Onishi (Duke University), Steve Haase (Duke University), and members of the Lew lab for stimulating discussions and comments on the manuscript. This work was funded by NIH/NIGMS grant R35GM122488 to DJL.

## AUTHOR CONTRIBUTIONS

Conceptualization, drafting, review, and editing manuscript— A.V. Colarusso, A.C.E. Wirshing, and D.J. Lew. Data curation, investigation, methodology, visualization, validation, and formal analysis— A.V. Colarusso and A.C.E. Wirshing. Project administration, supervision, and resources— D.J. Lew

## SUPPLEMENTAL MATERIAL

### SUPPLEMENTAL FIGURE LEGENDS

**Figure S1:**
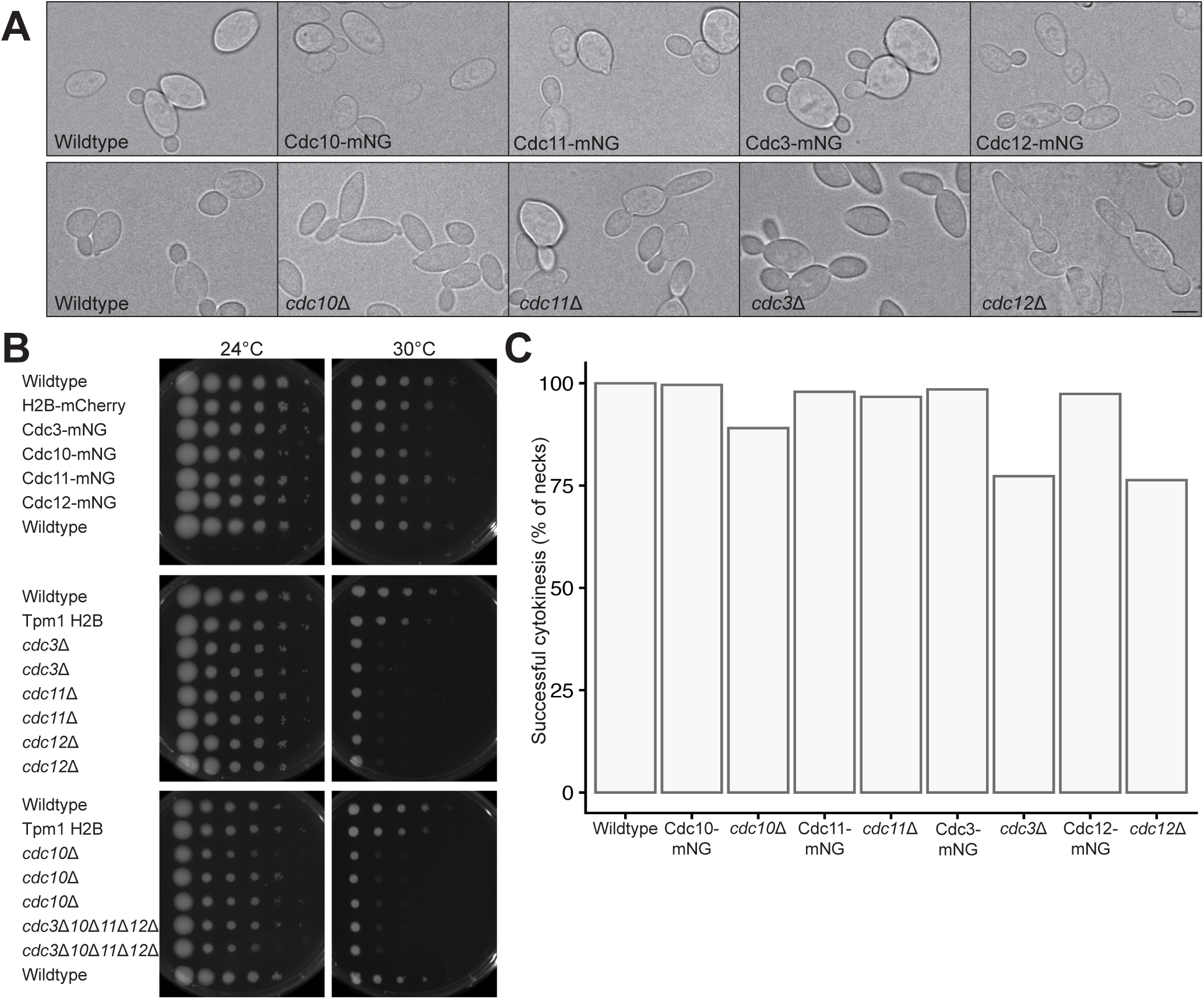
Growth of strains with septin tags and deletions. (A) Brightfield images of cells with the indicated septin-mNG tags (top row) or septin deletions (bottom row). Scale bar, 5 μm. (B) Colony growth of 10-fold serial dilutions of the indicated strains grown for 2 days on YPD at the indicated temperature. Septin deletes are severely temperature sensitive, which is largely but not completely complemented by the fluorescently tagged septins. (C) Percent of bud necks that complete cytokinesis in each tagged strain and deletion scored using brightfield imaging (n = 271, 248, 219, 239, 240, 167, 207, 230, 207 necks for wildtype, Cdc10-mNG, *cdc10*Δ, Cdc11-mNG, *cdc11*Δ, Cdc3-mNG, *cdc3*Δ, Cdc12-mNG, and *cdc12*Δ). See Table I for strains.

**Figure S2:**
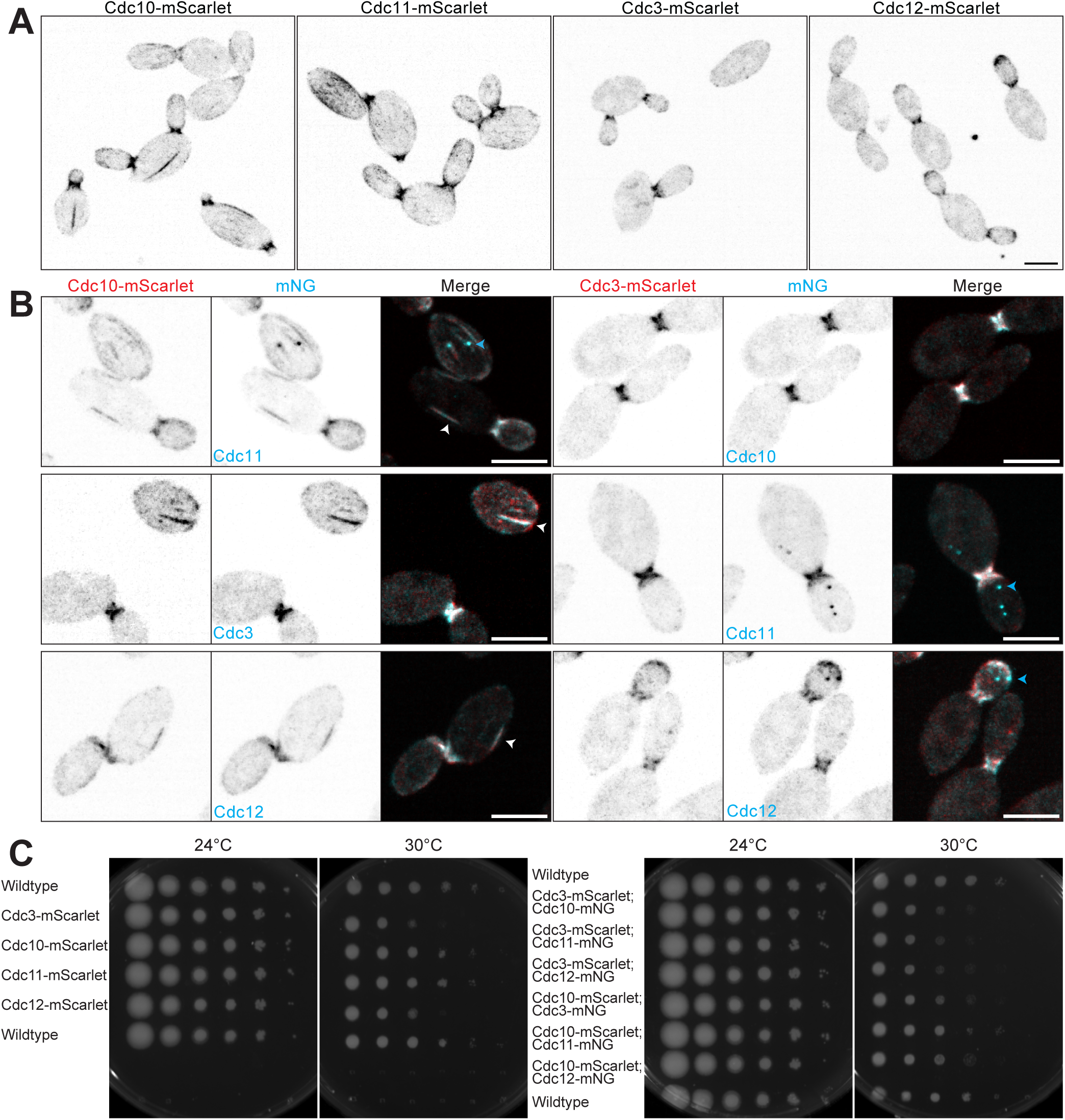
Two-color septin imaging. (A) Maximum projection confocal images of budding cells with the indicated septins tagged with mScarlet (Cdc10-mScarlet, DLY27419; Cdc11-mScarlet, DLY27423; Cdc3-mScarlet, DLY27416; Cdc12-mScarlet, DLY27426). Scale bar, 5 μm. (B) Maximum projection confocal images of cells expressing either Cdc3 or Cdc10 tagged with mScarlet (red) and a second septin tagged with mNG (cyan) (Cdc3-mScarlet Cdc10-mNG, DLY27568; Cdc3-mScarlet Cdc11-mNG, DLY27571; Cdc3-mScarlet Cdc12-mNG, DLY27575; Cdc10-mScarlet Cdc11-mNG, DLY27581; Cdc10-mScarlet Cdc3-mNG, DLY27578; Cdc10-mScarlet Cdc12-mNG, DLY27583). Cortical bars were all labeled with both tagged septins (white in overlay: white arrowheads). Cytoplasmic puncta were often labeled with only one tagged septin (cyan arrowheads). Scale bar, 5 μm. (C) Colony growth of 10-fold serial dilutions of the indicated strains grown for 2 days on YPD at the indicated temperature.

**Figure S3:**
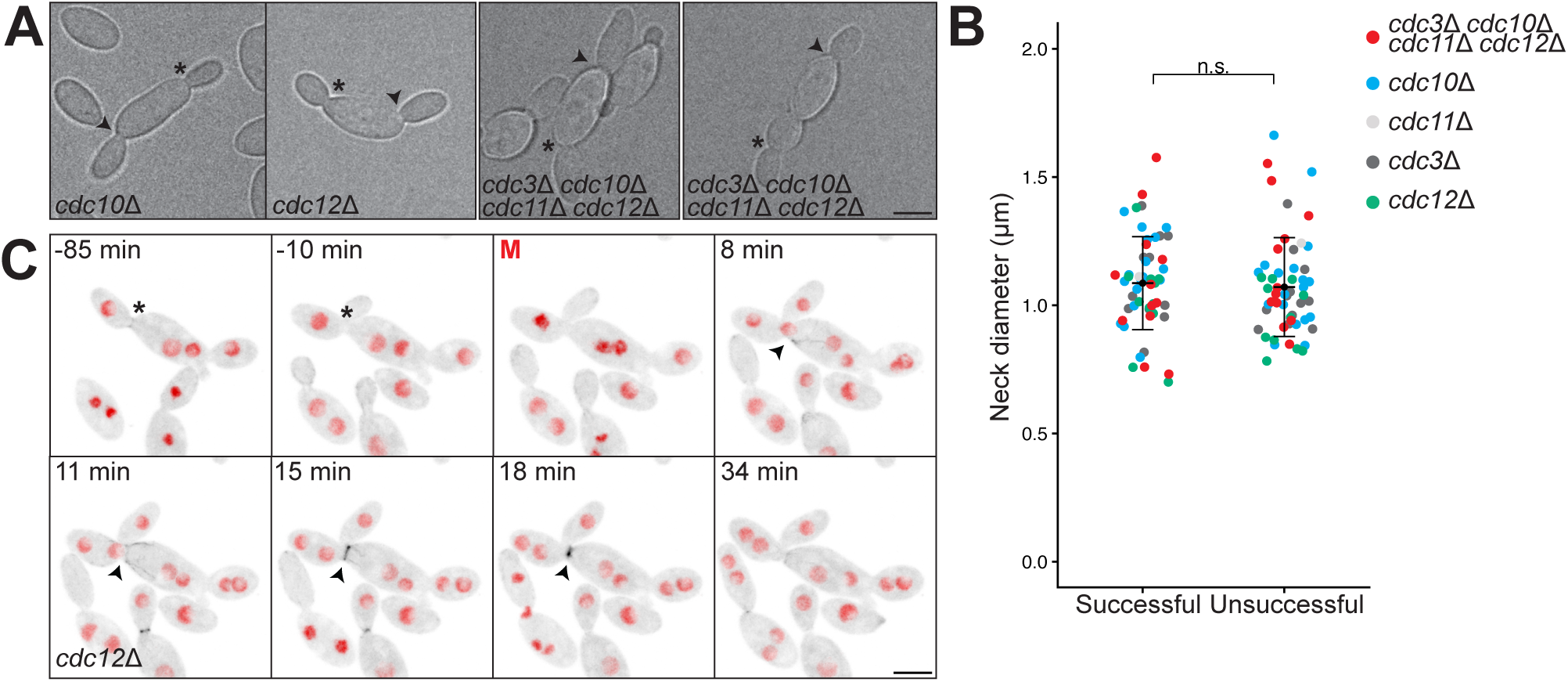
Cytokinesis success and neck diameter in septin mutants. (A) Brightfield images of two-budded cells of the indicated septin mutants at the time of mitosis. Arrowheads indicate bud necks that underwent successful cytokinesis, and asterisks indicate bud necks that failed to undergo cytokinesis. Strains: *cdc10*Δ, DLY26180; *cdc12*Δ, DLY26185; *cdc3*Δ *cdc10*Δ *cdc11*Δ *cdc12*Δ, DLY27187. Scale bar, 5 μm. (B) Bud neck diameter in the indicated mutants (color) at necks that did (successful) or did not (unsuccessful) undergo cytokinesis. Mean and standard deviation (n = 45 cells). Statistical significance calculated by a paired t-test (n.s., p > 0.05). (C) Time-lapse maximum intensity projection confocal images of actin (mNG-Tpm1, black) and nuclei (H2B-mCherry, red) in a *cdc12*Δ (DLY26186) cell that completes cytokinesis at a previously unsuccessful bud neck. Scale bar, 5 μm.

**Figure S4:**
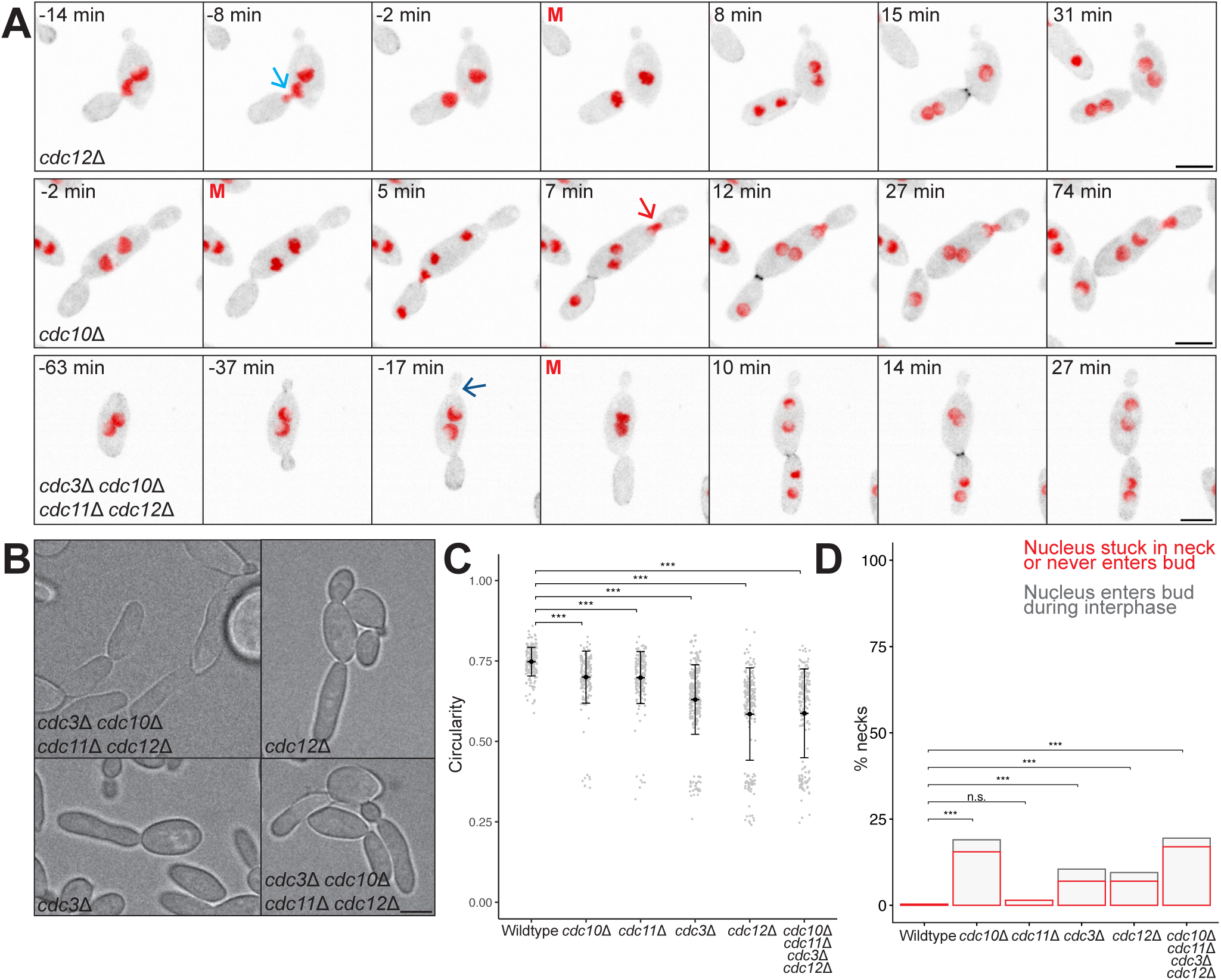
Septin mutant phenotypes beyond cytokinesis. (A) Time-lapse maximum intensity projection confocal images of actin (mNG-Tpm1, black) and nuclei (H2B-mCherry, red) in the indicated septin mutants (*cdc12*Δ, DLY26186; *cdc10*Δ, DLY26180; *cdc3*Δ *cdc10*Δ *cdc11*Δ *cdc12*Δ, DLY27187). Cyan arrow indicates nucleus entering bud during interphase, red arrow indicates a mitotic nucleus stuck in the bud neck, and dark blue arrow indicates a bud that stops growing prematurely. Scale bar, 5 μm. (B) Brightfield images of cells of the indicated mutants that developed elongated buds (*cdc12*Δ, DLY26185; *cdc3*Δ, DLY26176; *cdc3*Δ *cdc10*Δ *cdc11*Δ *cdc12*Δ, DLY27187 or DLY27188). Scale bar, 5 μm. (C) Circularity of buds for the indicated mutants with mean and standard deviation shown (n = 200 buds each). Statistical significance calculated by pairwise Wilcoxon test (n.s., p > 0.05; ***, p ≤ 0.005). (D) Percent of buds displaying nuclear errors (n = 200 buds each). Statistical significance calculated by Chi-squared test (n.s., p > 0.05; ***, p ≤ 0.005).

**Figure S5:**
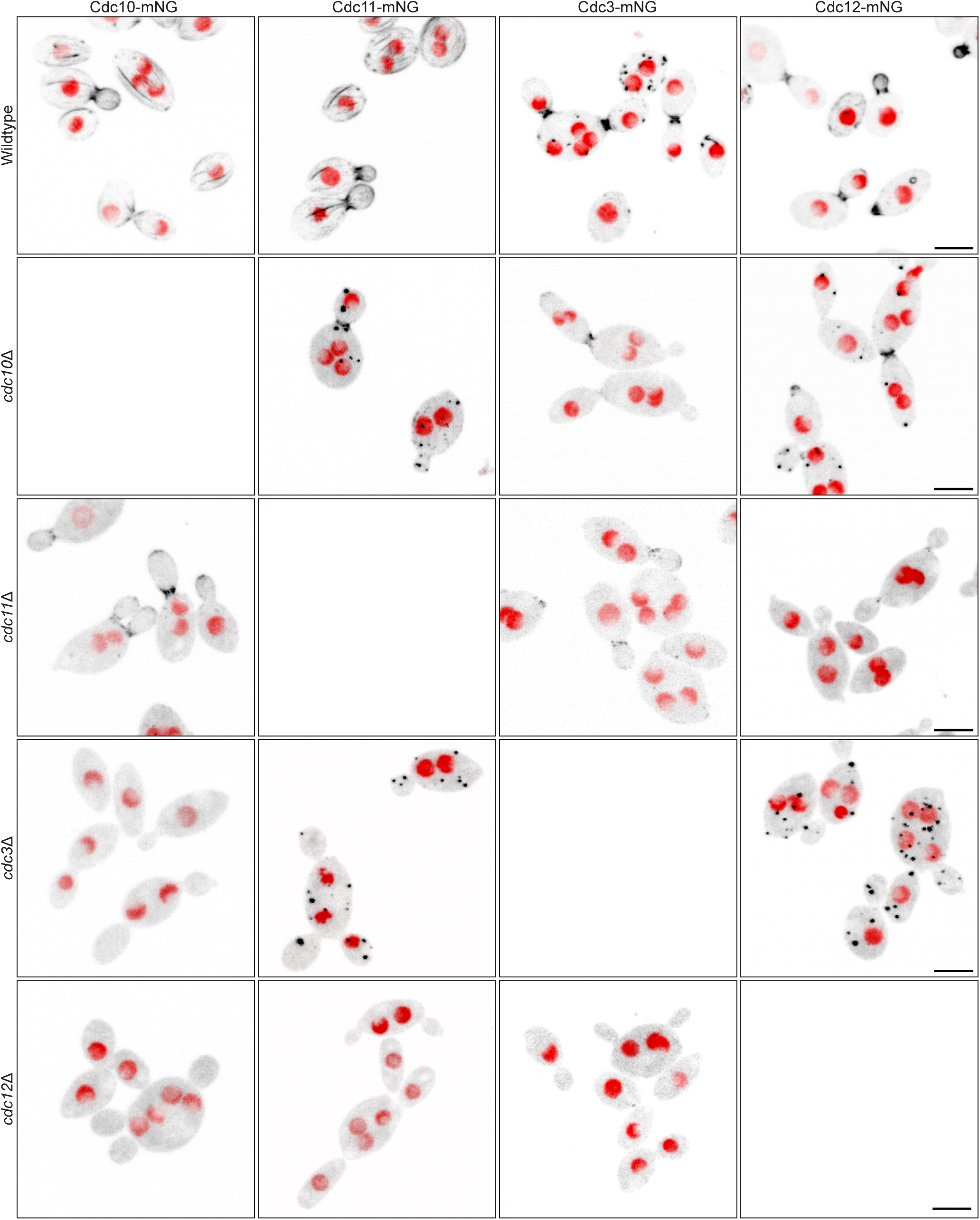
Septin localization in the absence of other septins. Merged maximum intensity projection confocal images of cells expressing H2B-mCherry (red) and the indicated septin-mNG (columns) in the indicated strains (rows). See Table I for strains. Scale bar, 5 μm.

### SUPPLEMENTAL VIDEO CAPTIONS

**Video 1: Actin and nuclei in *A. pullulans* yeast**

Actin visualized with mNG-Tpm1 (black) and nuclei with H2B-mCherry (red). Merged maximum projection time lapse images of DLY26075 or DLY26076 at 1 min intervals. Scale bar, 5 μm.

**Video 2: Actin and nuclei in *A. pullulans* hyphae**

Actin visualized with mNG-Tpm1 (black) and nuclei with H2B-mCherry (red). Merged maximum projection time lapse images of DLY26075 at 2 min intervals. Scale bar, 10 μm.

**Video 3: Septins and nuclei in *A. pullulans* yeast**

Merged maximum projection time lapse images of Cdc3-mNG (DLY26842), Cdc10-mNG (DLY26846), Cdc11-mNG (DLY26848), and Cdc12-mNG (DLY26802) at 1 min intervals. Scale bar, 5 μm.

**Video 4: Septins and nuclei in *A. pullulans* hyphae**

Merged maximum projection time lapse images of Cdc3-mNG (DLY26842), Cdc10-mNG (DLY26846), Cdc11-mNG (DLY26848), and Cdc12-mNG (DLY26802) at 2 min intervals. Scale bar, 10 μm.

**Video 5: Cytokinesis in septin mutant yeast cells**

Actin visualized with mNG-Tpm1 (black) and nuclei with H2B-mCherry (red). Merged maximum projection time lapse images of *cdc12*Δ (DLY26185), *cdc10*Δ (DLY26180), *cdc3*Δ *cdc10*Δ *cdc11*Δ *cdc12*Δ (DLY27188), and *cdc3*Δ (DLY27177) mutants at 1 min intervals. Scale bar, 5 μm.

**Video 6: Cytokinesis in septin mutant hyphae**

Actin visualized with mNG-Tpm1 (black) and nuclei with H2B-mCherry (red). Merged maximum projection time lapse images of wildtype (DLY26075) and *cdc3*Δ *cdc10*Δ *cdc11*Δ *cdc12*Δ (DLY27187) mutants at 2 min intervals. Scale bar, 5 μm.

## ABBREVIATIONS

CAR: contractile actomyosin ring
Hyg: hygromycin B
mNG: mNeonGreen
Nat: nourseothricin

